# Electrical Fields Induced Inside the Rat Brain with Skin, Skull, and Dural Placements of the Current Injection Electrode

**DOI:** 10.1101/402578

**Authors:** Ahmet S. Asan, Sinan Gok, Mesut Sahin

**Affiliations:** Department of Biomedical Engineering, New Jersey Institute of Technology, University Heights, Newark, NJ 07102 USA; Professor of Biomedical Engineering, Associate Editor, IEEEE Tran. on Biomedical Circuits and Systems, Department of Biomedical Engineering, New Jersey Institute of Technology, University Heights, Newark, NJ 07102 USA

**Keywords:** transcranial electrical stimulation, electrical field distribution, volume conductor

## Abstract

Transcranial electrical stimulation (tES) is rapidly becoming an indispensable clinical tool with its different forms. Animal data are crucially needed for better understanding of the underlying mechanisms of tES. For reproducibility of results in animal experiments, the electric fields (E-Fields) inside the brain parenchyma induced by the injected currents need to be predicted accurately. In this study, we measured the electrical fields in the rat brain perpendicular to the brain surface, i.e. vertical electric field (VE-field), when the stimulation electrode was placed over the skin, skull, or dura mater through a craniotomy hole. The E-field attenuation through the skin was a few times larger than that of the skull and the presence of skin substantially reduced the VE-field peak at the cortical surface near the electrode. The VE-field declined much quicker in the gray matter underneath the pial surface than it did in the white matter, and thus the large VE-fields were contained mostly in the gray matter. The transition at the gray/white matter border caused a significant peak in the VE-field, as well as at other local inhomogeneties. A conductivity value of 0.57 S/m is predicted as a global value for the whole brain by matching our VE-field measurements to the field profile given by analytical equations for volume conductors. Finally, insertion of the current return electrode into the shoulder, submandibular, and hind leg muscles had virtually no effects on the measured E-field amplitudes in the cortex underneath the epidural electrodes.

## I. INTRODUCTION

Transcranial electrical stimulation (tES) has emerged as an effective non-invasive technique for modulation of the brain activity in recent years, while the earliest studies date back more than a century (1). In general, the safety of tES and its derivations has now been agreed upon so long as the current is kept below 2mA, although recent reports seem to suggest 4mA as the limit (2). These tES derivations include transcranial direct current stimulation (tDCS), transcranial alternating current stimulation (tACS), and transcranial random noise stimulation (tRNS) (3,4). Despite widespread interest in clinical applications, the exact mechanisms of action for tES are still being investigated.

Aforementioned tES techniques inject weak electrical currents through the brain and presumably cause only subthreshold modulation of the neuronal membrane potentials (5). Early animal studies demonstrated that neuronal firing rates could be modulated by applying direct currents (DC) to the brain (6,7) and the modulatory effects of transcranial DC electric fields were confirmed in human studies (8,9). While the excitatory and inhibitory effects of tDCS are attributed to the direction of the applied current on a larger scale (anodal vs. cathodal) (7,8,10), a more detailed analysis revealed that the primary factor is the position and orientation of the individual neuronal structures relative to the electric field. *In vitro* measurements on rat hippocampal slices showed that the amplitude and delay of the population spikes evoked by orthodromic stimulation were affected linearly by the applied uniform electric fields (11). Furthermore, even small electric fields induced polarization of CA1 pyramidal cells of the hippocampus when applied parallel to the somato-dendritic axis, but failed to do so when applied perpendicularly. Studies on the rat motor cortex slices indicated that somatic polarization was also correlated with the neuronal morphology, and the layer V pyramidal cells were the most sensitive to subthreshold electric fields (12). In general it seems that a realistic estimation of the electric field distribution inside the brain parenchyma is needed as a starting point for accurate interpretations of the neurological impact of the intervention.

The electric field can be controlled by adjusting the stimulation current/charge intensity and steered to a certain extent by careful positioning of the extracranial electrodes. Nevertheless, the electrical properties of different tissues that the current passes through play a significant role in distribution of the electric field. Direct in vivo measurement of the electric field in human subjects is not an option from a clinical standpoint, thus most investigators resort to computational models in order to approximate the electric field distribution in the human brain under varying stimulation intensities and electrode arrangements. Miranda et al. used a spherical head model and estimated that almost 50% of the injected current is shunted through the scalp (13). Datta et al. used a more advanced head model where gyri/sulci specificity was defined and predicted that the electric field was concentrated at distinct sites, like the walls of the gyri (14). In summary, tES induced current flow is influenced by several factors including: 1) skull and scalp thicknesses and their compositions, 2) cerebrospinal fluid (CSF) thickness and conductivity, 3) gyri/sulci morphology, 4) electrode size and geometry, and 5) positioning of the stimulating and return electrodes (14,15).

In addition to human computational models, tES effects have been studied in animals where the main interests are understanding the underlying cellular and molecular mechanisms, optimizing stimulation protocols, and establishing safety limits. To this end, animal models provide ample opportunities to rapidly develop new tDCS methodologies and measure the outcomes while manipulating the stimulation parameters within a large range that may not be feasible clinically (see (16) for a review). For accurate interpretation of the results from these animal studies, realistic estimates of the induced electric field distribution are needed. Direct measurement of the electric fields in brain tissue goes back to as early as 1950s with experiments carried on anesthetized monkeys (17), although not many follow-up studies reported since then. Chan et al. conducted a series of experiments using isolated turtle cerebella, and studied the relationship between the applied fields and the spontaneous neuronal activity (18,19). More recent studies on monkeys (20) and rats (21) also reported electric field measurements, although restricted either to the cortical surface or a single horizontal plane, respectively.

The tDCS technique can benefit from in vivo animal data, which are clearly lacking in the literature, for better understanding of the mechanisms. While the computational models provide a basic understanding of how electrical currents are distributed through the animal brain (22) as much as the human brain, they can lead to unrealistic conclusions if they primarily depend on conductivity measurements gathered from *ex vivo* tissue samples (23). Furthermore, anatomical differences in the brain size and the skin, skull, and CSF thicknesses make it difficult to extrapolate the results of an electric field model developed for one species to another. In this study, we used a rat model to measure the intracerebral voltages at varying depths and horizontal distances from the stimulating electrode. Vertical electric fields were reported for three stimulation conditions in which the anodic electrode was placed over the shaved skin, the skull, and the dura mater. The rat brain was selected in this study due to its common usage as an animal model. We anticipate that the results of this study will provide a reference point for more realistic estimations of the electric field distribution in other studies using the rat model and help to improve the reproducibility of the reported tES effects. Some of the results, such as the shunting effect of the skin and the CSF, agree with modeling predictions, while some others such as the relatively smaller attenuation by the rat skull, the E-field peaks at the white/gray matter border, and insensitivity of the field to the reference electrode location are among the practical findings of this study.

## II METHODS

### Animal Surgery

Ten Sprague Dawley rats (250-350g, male) were used in this study for direct measurements of the electric field distribution in the brain parenchyma. This study was carried out in strict accordance with the recommendations in the Guide for the Care and Use of Laboratory Animals of the National Institutes of Health. The protocol was approved by the Institutional Animal Care and Use Committee (IACUC), Rutgers University, Newark, NJ (Protocol Number: 201702616). Anesthesia was induced with 5% isoflurane gas in an induction chamber, maintained by 1-3% isoflurane in 95% oxygen after moving the animals to a stereotaxic frame, and monitored using the toe pinch reflex. All efforts were made to minimize suffering. Blood oxygen level was monitored via a pulse oximeter (NPB-40, Nellcor Puritan Bennet) from the hind paw. Body temperature was measured with a rectal temperature probe (World Precision Instruments-WPI) and regulated with a heating pad (WPI) underneath the animal over the course of surgery. The hair over the head was shaved with an electric shaver and the skin was treated with a depilatory cream to remove the fine hair. The skin was then cleansed with antiseptic solution.

### Current Injection over the Skin

Either 4mm diameter pellet electrodes (EP4, Ag/AgCl, WPI, FL) or 1.5mm diameter helical wire electrodes were used to inject the electric currents. The pellet electrodes are commercially available and thus more common in tES experiments in animals, whereas the helical electrode that is hollow in the center allowed us to make E-field measurements underneath the electrode. The pellet electrode was positioned such that medial edge was juxtaposed to the sagittal suture and the caudal edge to the coronal suture. For helical electrodes, a Ag/AgCl wire with 125µm uncoated thickness (A-M Systems, #786000) was wrapped around a 1.25mm diameter rod 4 times to form a helix with a large surface area and thus a lower impedance. The impedance was confirmed to be below 10kΩ @1kHz in phosphate buffered saline (PBS, Sigma-Aldrich). The helical electrode was filled with conductive gel in the center to ensure good contact with the skin and distribute the current more uniformly at the base of the electrode. The helical electrode was placed with its center positioned 2mm lateral (left or right) from the sagittal suture and 2mm either rostral or caudal to the coronal suture. Another Ag/AgCl wire was inserted to the ipsilateral shoulder muscles as the return (cathodic) electrode for the injected current. Ten monophasic anodic pulses were delivered as a train at 100µA amplitude, 100ms pulse width, and a repetition rate of 4Hz. The amplitude was switched to 200µA at times to increase the signal-to-noise ratio where the recorded amplitudes were too small. In this protocol, substituting pulsed stimulation for DC allowed us to overcome the poor DC response of the metal recording electrodes.

A sharp cut was made into the skin with a surgical blade at the edge of the stimulation electrode to expose the skull caudally while leaving the skin underneath the electrode intact (Fig. 1B). Two 1mm diameter craniotomy holes were drilled 2mm and 4mm away from the caudal edge of the stimulation electrode using a micro drill (OmniDrill 35, WPI). A tungsten electrode (0.5 MΩ, TM33B05H, WPI) was inserted into the craniotomy hole to record the induced voltages as a function of depth with respect to another Ag/AgCl reference electrode attached on the skull near the recording electrode using dental acrylic. The dura was punctured with the sharp tip of the tungsten electrode and the first recording was made at the level of the cortical surface (depth=0). Using a 10µ-resolution micromanipulator (Kite-R, WPI), the tungsten electrode was advanced into the brain parenchyma in 0.2mm steps until reaching 4mm depth and thereafter in 0.5mm steps up to a depth of 6mm. The procedure was repeated in both craniotomy holes, at 2mm and 4mm horizontal distances from the stimulation electrode. The rising and falling edges of the recorded signals were marked using an automated algorithm in MATLAB (Mathworks Inc.) to quantify the induced voltage amplitudes (see Fig. 2 inset). In some animals, this procedure was repeated on the contralateral side of the brain to obtain an additional set of measurements.

**Fig. 1:**
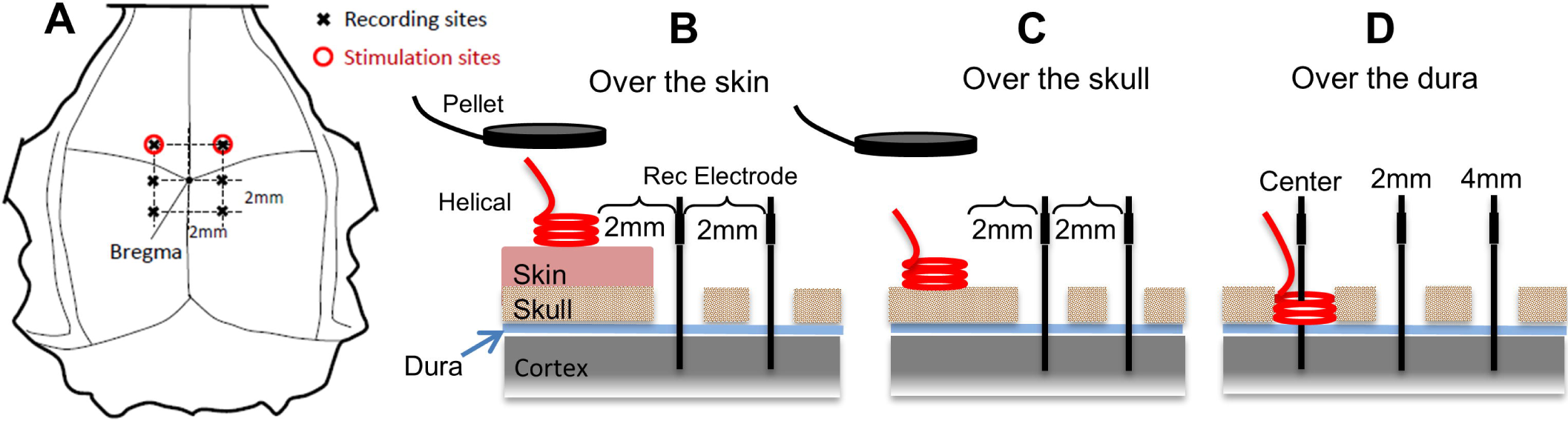
Drawings show the three different placements of the stimulation electrode on the rat’s head: A) top view. The vertical cross sections of the skin and skull in B, C, and D show the placements of the helical wire stimulation electrode: B) over-the-skin, C) over-the-skull, and D) over-the-dura, and the craniotomy holes for E-field measurements. Tungsten recording electrodes were inserted through the craniotomy holes at center (for over-the-dura stimulation only), and 2mm and 4mm horizontal distances from the stimulation electrode. The pellet electrodes were tested only over-the-skin and skull as shown in B and C.

**Fig. 2:**
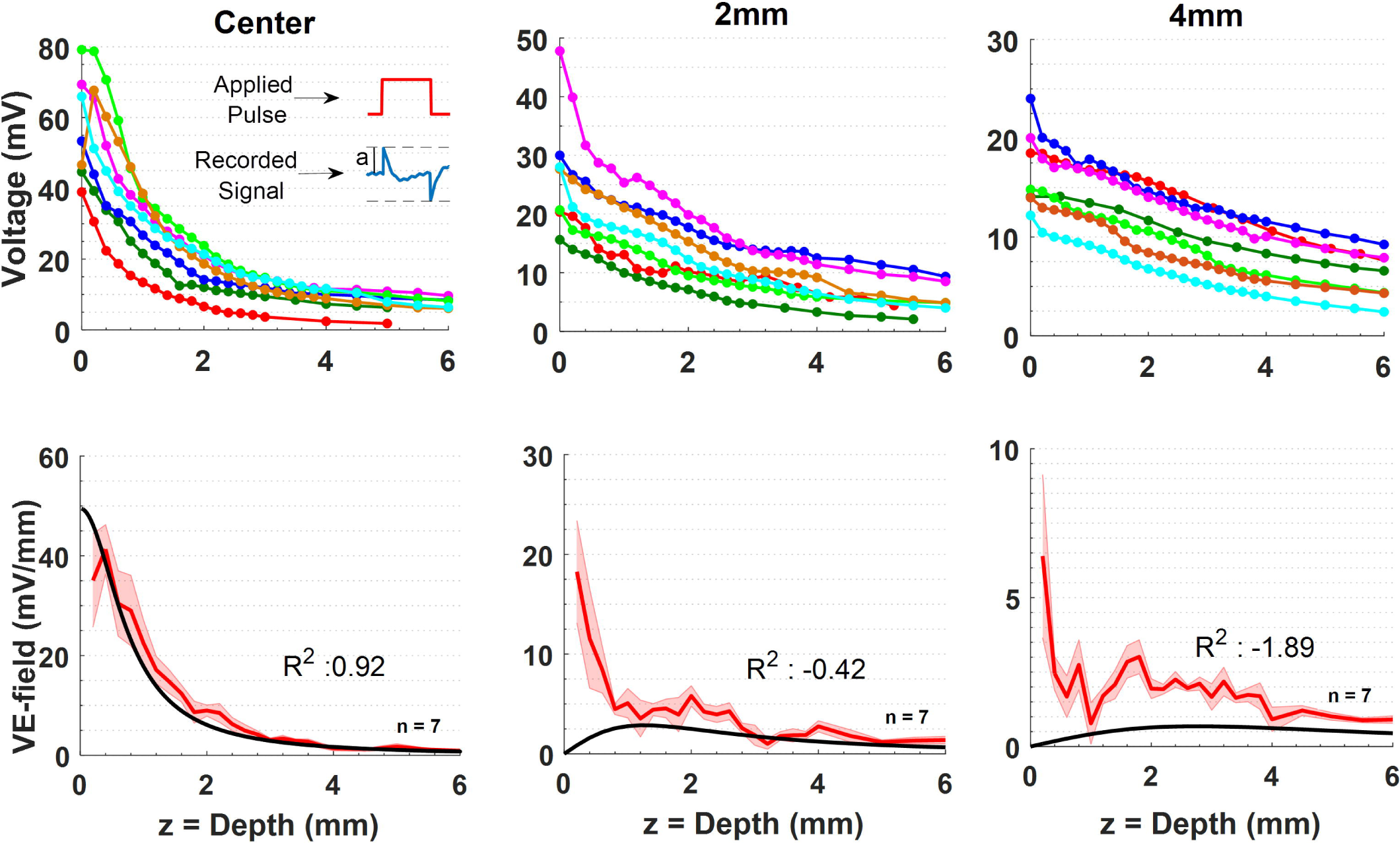
Voltage (top row) and VE-field (bottom row) measurements made with epidural (as in Fig. 1D) placement of the Ag/AgCl helical wire stimulation electrode with 1.5 diam. Top Row Left to Right: Voltage measurements made with respect to a reference on the skull and via penetrations at the center, and 2mm and 4mm from the edge of the helical electrode. Seven sets of measurements were collected in five animals. In two sets, the voltage was measured at a fewer points below 3mm of depth. In such cases, exponential interpolation was utilized to estimate the missing voltage values. The inset depicts how the voltage amplitude (a) measurements were made at the rising edge of the recorded waveforms. Bottom Row: The mean of the vertical E-fields calculated by differentiating the voltage measurements shown in the top row for each penetration separately. The shaded areas indicate ± standard error. The black solid line is the VE-field predicted by the analytical equation derived by Wiley & Webster (24) that provided the best fit, *i.e*. highest coefficient of determination (R2), when Vo at V(r=0, z=0) is set to 58.3mV. For 2mm and 4mm penetrations, the analytical equation was evaluated (solid black lines) using the same Vo value for consistency.

### Current Injection over the Skull

The skin over the top of the skull was completely removed in this step, mostly in the same animals used above. After removing the periosteum, bone wax was applied to the muscles around the edges and the skull sutures on top to stop bleeding. The stimulation electrode (pellet or helical wire) was placed onto the skull (Fig. 1C) at the same coordinates used with over-the-skin electrodes measured with respect to the *bregma*. The voltage measurements were made following the same procedure above with slight repositioning of the tungsten electrode in the same craniotomy holes in order to avoid damaged tissue from the previous penetration.

### Current Injection over the Dura

A 2mm craniotomy hole was made at the coordinates of the stimulation electrode to inject the current directly through the dura (Fig. 1D). A helical wire electrode (1.5mm diam.) was placed about 0.5mm above the dura mater and anchored to the edge of the skull hole with small amounts of cyanoacrylate glue and the hole was filled with normal saline. The distances between the caudal edge of the helical electrode and recording holes were kept at 2mm and 4mm as before. With the helical electrodes, an additional set of voltage recordings were made by inserting the tungsten electrode through the center of the stimulation electrode.

Finally, in order to investigate the effect of the current return electrode position, the Ag/AgCl wire inserted to the ipsilateral shoulder muscles was moved to the contralateral shoulder, the hind leg, and the submandibular muscles via needle insertions. The voltage measurements were repeated near the cortex and at various depths with the recording electrode in the center of the helical electrode for these four different positioning of the current return electrode for comparison.

### Data Collection and Analysis

The signals were collected in a large Faraday cage and first amplified by a gain of 100 (Model 1700, A-M Systems, WA) with filters setting of 10 Hz–10 kHz, and then sampled at 25kHz through a National Instruments data acquisition board (PCI 6071) controlled by custom-designed MATLAB codes. Stimulus-triggered averaging (STA) was used (N=10) to suppress the background neural activity and other sources of random noise. The averaged signal was further band-pass filtered in MATLAB (10 Hz – 1 kHz) before analyzing. Voltage transitions within 2ms window around the rising edge of the square pulses were taken as the induced voltage at the corresponding depth. The first derivative of the voltage data with respect to depth was computed as the electric field (E-field). All field measurements were made exclusively in the vertical direction (VE-field) in this study. Horizontal E-field measurements would require derivation of the voltage measurements made through different brain penetrations and lead to large calculation errors due to even slight changes in the absolute value of the voltages measured, which could easily occur from repositioning of the recording reference electrode between penetrations.

### Comparison with Theoretical Models

The resistance of a monopolar disk electrode at the surface of a semi-infinite, homogeneous, and isotropic medium was first derived by Newman (24) as a function of the electrode radius (a), and the conductivity of the medium (σ). The potential at the surface is found as *Vo = J/(4σa)*, where I is the electrode current. Wiley and Webster then solved the Laplace’s equation for the voltage distribution inside the semi-infinite volume conductor for a similar disk electrode (25). By substituting zero for the horizontal axis r in their equation, we find the voltage profile at the electrode center, which starts from Vo at z=0 and declines with increasing values of the vertical axis (z) according to the equation 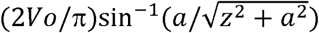. The VE-field, however, decreases as a function of *ϕ = –(2V*_*o*_*/π)(a/z*^*2*^ *+ a*^*2*^), which can be found easily by differentiating the voltage equation with respect to z, and the peak value of the VE-field at the surface (z=0, r=0) is *ϕ = –2V*_*o*_*/(πa)* or *ϕ = J/(2πσa*^*2*^). We fit the VE-field equation to our data collected with the epidural placement of the electrode at its center while leaving Vo as a free parameter (Fig. 2, center). The Vo value that fits the experimental data best for r=0 was also used to plot the VE-field equation at r=2mm and r=4mm from the edge of the stimulation electrode in Fig. 2 (2mm and 4mm). The VE-field is shown as a 2D heat plot in Fig. 3 for comparison with the experimental data and will be discussed at the end.

**Fig. 3:**
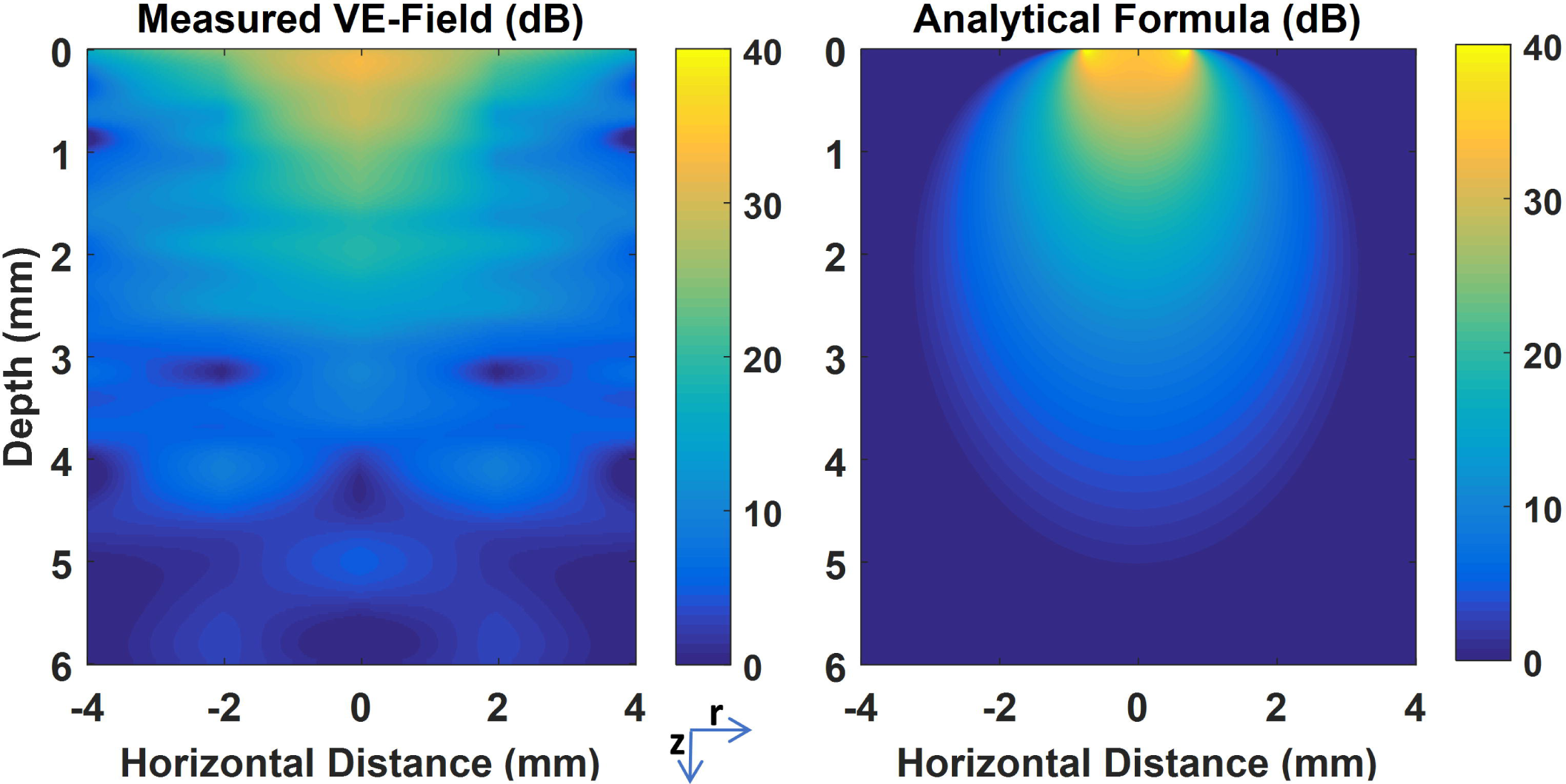
Vertical E-field distribution as a heat plot on a logarithmic scale. The left panel was obtained by linear interpolation of the measurements presented in Fig. 2 and reflected over the vertical axis in the middle. The right panel is the predicted VE-field by the analytical equations of Wiley & Webster. Depth=0 corresponds to the surface of the cortex.

## III RESULTS

### VE-Fields with Epidural Injection of Current

The peak-to-peak voltage induced by the current pulse was measured as a function of depth down to 6mm from the dura surface (Fig. 2, top row) at the center of the helix, and 2mm and 4mm away horizontally from the stimulation electrode (see Fig. 1D). Vertical electrical fields (VE-fields) were computed by differentiating the raw voltage measurements without any filtering or curve fitting, and averaged across the animals (Fig. 2, bottom row). Note that the vertical offset in the voltage plots of the top row do not carry any significance since they can vary between animals depending on where the recording reference electrode is placed along the current pathways between the stimulation and the return electrodes. The voltage curves decline sharply under the stimulation electrode with maximum values underneath the dura (depth=0mm), as expected. The VE-field was also maximum under the stimulation electrode (center, depth=0mm) near the cortex, declined exponentially, and lost more than 75% of its strength by 2mm below the stimulation electrode and further decreased down to negligible levels by 6mm. The analytical equation by Wiley and Webster (see methods) was evaluated and fit to the VE-field by using Vo as the free parameter. The analytical formula provided a good-fit (R =0.92) including the initial plateau near the cortical surface for Vo=58.3mV. As the recording electrode was moved horizontally to 2mm and 4mm away from the stimulation site, the VE-field decline near the surface became sharper and an elevation appeared at deeper levels. The initial sharp decline was not predicted by the analytical equation since the semi-infinite model assumes a non-conductive medium above the surface and a zero vertical current at the boundary. The band of large VE-fields extending horizontally underneath the surface can also be appreciated from the heat plot of Fig. 3A. Interestingly, for all penetrations (center, 2, 4mm) there seems to be a peak in the E-field near or a little above the 2mm mark, which may indicate a sharp change in tissue conductivity, e.g. from gray matter to white matter. The analytical equations fail to predict the VE-field amplitudes, in general, at 2mm and 4mm from the stimulation electrode with the same Vo used at center penetration. Thus, the VE-fields expand more in the horizontal direction than predicted.

### Comparison of VE-Fields with Current Injections over the Skin, Skull, and Dura

Figure 4 compares the VE-fields with three different placements of the helical electrode; over the skin, skull, and dura. The epidural stimulation produces the largest electric field intensities in the brain parenchyma while epidermal placement produces the lowest intensity, as expected, both at 2mm and 4mm horizontal locations from the electrode. Note that for over-the-skin and skull placements of the stimulation electrode the E-fields measurements were not in the center of the electrode in order not to disturb the intactness of the skin or skull with a penetration hole. The skin attenuated the VE-field (or shunted the electric currents) to a much larger degree than the skull, as seen with penetrations both at 2mm and 4mm horizontal locations from the stimulation electrode. The over-the-skin placement did not produce an exponentially decreasing VE-field profile by depth as the other two placements of the stimulation electrode, and the VE-fields measured at the cortical surface (depth=0) were about an order of magnitude smaller. The skin thickness was measured at the end of the experiment and found to be around 0.5mm under the stimulation electrode.

**Fig. 4:**
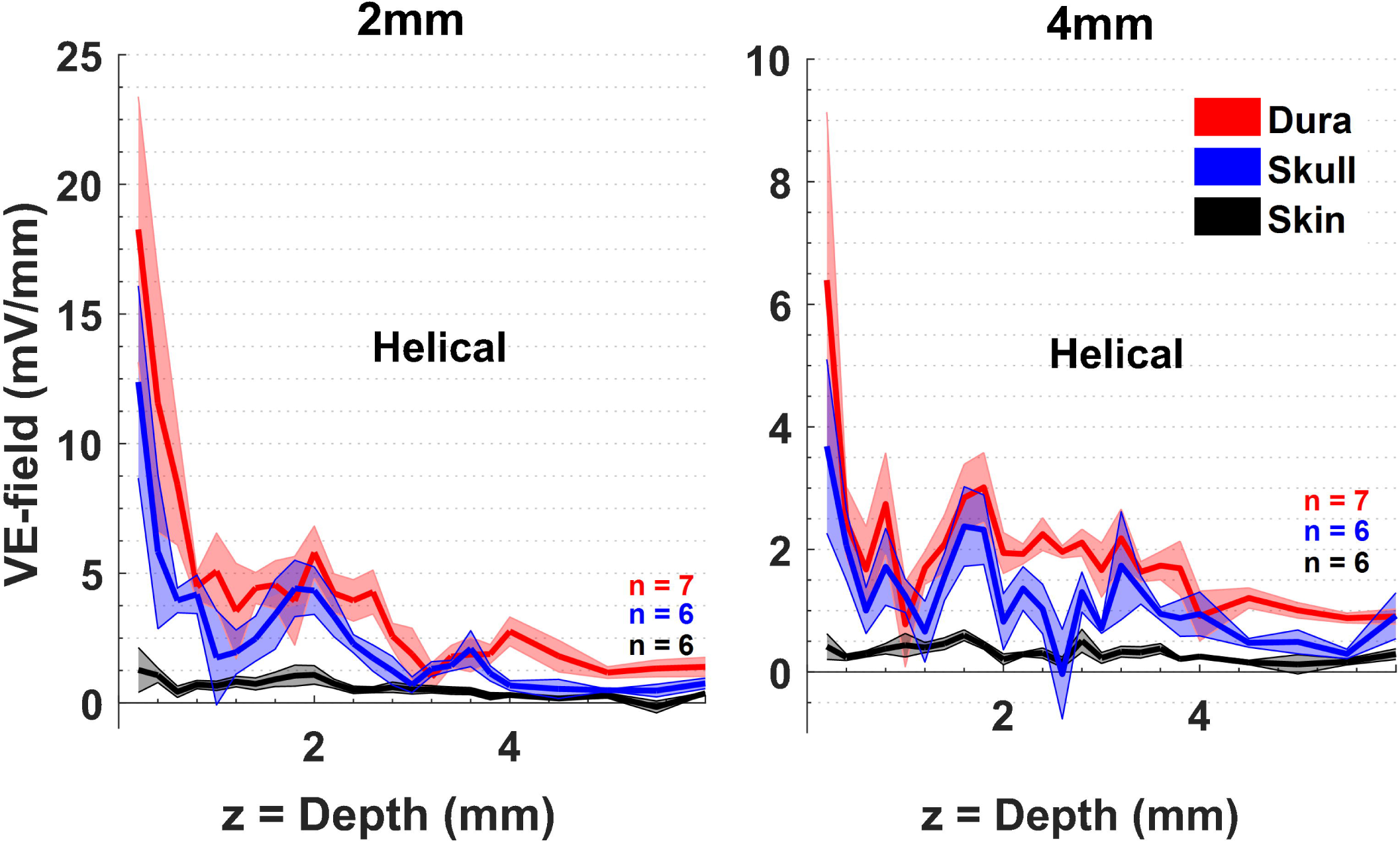
Comparison of VE-fields for three different placements of the helical wire electrode over the skin, skull, and dura mater. Measurements are repeated at 2mm (left) and 4mm (right) horizontal distances from the edge of the helical electrode. The averages of (n) measurement sets are plotted (solid lines) and the standard errors are shown as shaded areas.

The VE-fields were also compared with larger Ag/AgCl pellet electrodes (diam.=4mm) placed either over the skin or skull (Fig. 5). At a horizontal distance of 2mm from the edge of the pellet electrode, the presence of skin attenuated the electric fields by 3-4 times, similar to the smaller helical electrodes, but the VE-field decline was sharper by depth with the pellet electrode compared to the smaller helical electrodes. This was probably because the pellet electrode had a larger diameter and the 2mm from the edge was actually further away from the electrode center compared to the helical electrodes. That the VE-field decline within the first mm of depth becoming sharper as the penetrations move horizontally was also observed with the helical electrode (Fig. 2). The VE-field bump around 1.5- 2mm depth was observed consistently in these plots as well. For both skin and skull placements of the stimulation electrode, the measurements at 4mm were about a few times smaller than those at 2mm.

**Fig. 5:**
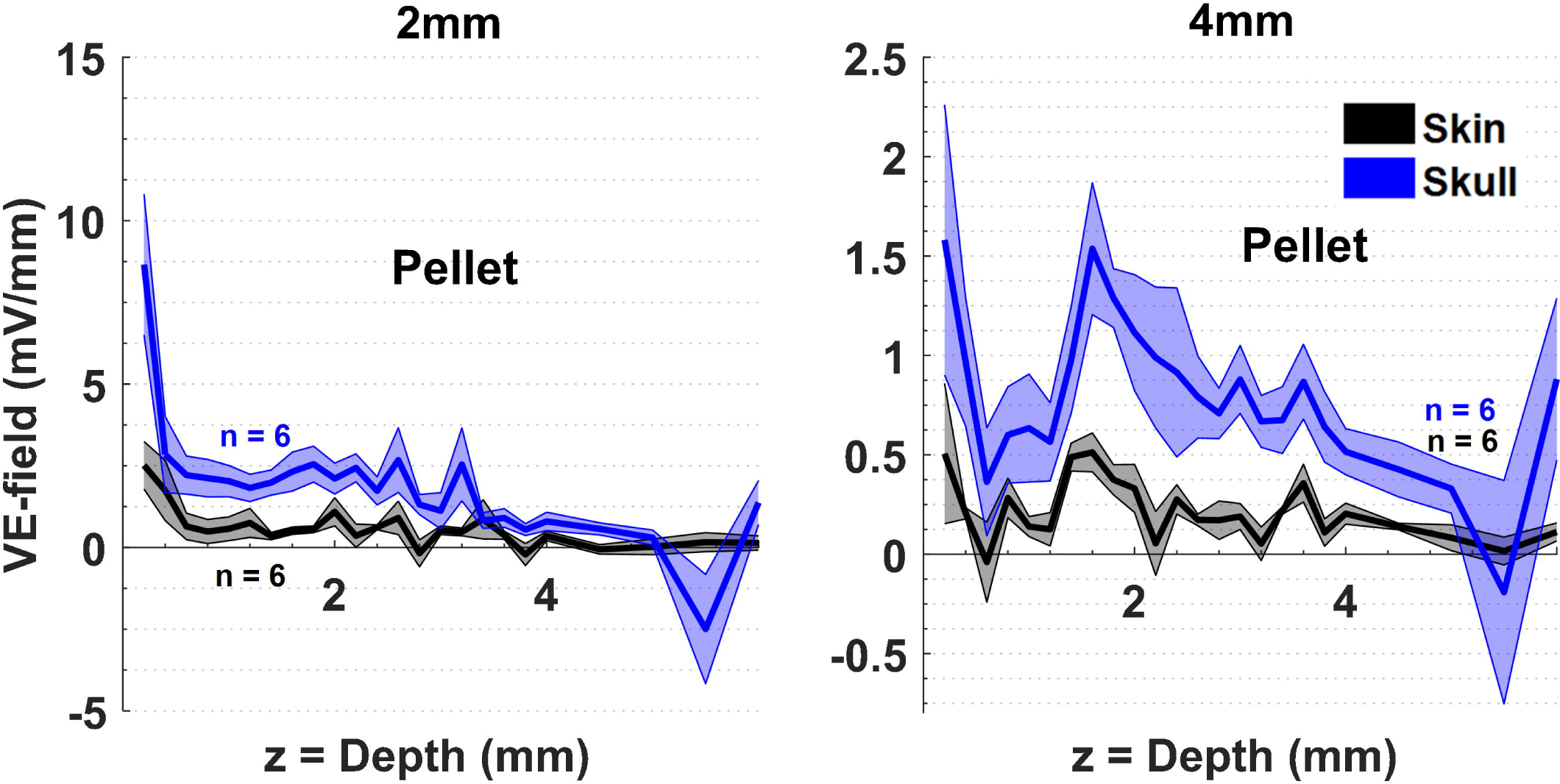
Comparison of VE-fields for two different placements of the large pellet electrode; over the skin and skull. Details are same as in Fig. 4.

### Location of the Current Return Electrode

Lastly, we investigated how the location of the stimulation reference electrode affects the electric field strength in two rats. In the previous experiments, a Ag/AgCl wire inserted into the ipsilateral shoulder was used as the current return electrode (cathode). For this experiment, three alternative sites were tested for the cathodic electrode: the contralateral shoulder, the submandibular muscles, and the ipsilateral hind limb. The epidural stimulation experiments with the helical wire electrodes were repeated against these reference electrode placements and the VE-fields as the difference of the voltage between the depths of 0.6-2.0mm were measured in nine different craniotomy holes (4 and 5 holes in two rats) as the stimulus was applied through the same hole. The maximum deviation from the ipsi-shoulder measurement was less than 1.36% with any of the new reference points. None of the VE-field measurement sets made at various depths from 0.6 to 2mm (N=5) against the novel reference points were significantly different from that of the set against the ipsi-shoulder electrode (paired t-test, p>0.5).

## IV DISCUSSION

### E-field Attenuation by Skin and Skull

The skin thickness changes around the head and rodents are no exception to this rule. Because of the compression we applied to hold the electrode down firmly on the skin, the skin thickness decreased under the stimulation electrode during the course of the experiment and found to be ∼0.5mm at the end. The skin had to be removed also caudal to the stimulation electrode to make craniotomy holes at the recording points. Hence, we can only make general remarks about the skin effects on the E-field measurements in the face of these sources of variability and practical limitations. About a four-fold decrease occurred in the VE-field when the electrode was placed over the skin as opposed to the skull surface. The average skull thickness was measured to be ∼0.5mm (N=5). In agreement with this data, Vöröslakos et al. reported that transcutaneous stimulation generated several-fold weaker electric fields compared to subcutaneous stimulation in rats (21). There was a relatively small loss of electric field intensity when the helical electrode was moved from the dural surface to the skull surface in our data.

With over-the-skin electrodes and 100µA current injection, the vertical electric field drops down to ∼1mV/mm at 2mm and 4mm from the stimulation electrode (Fig. 4). Terzulo and Bullock reported that the firing frequency of neurons can be modulated by voltage gradients as low as 1mV/mm (10), which is also the lower bound indicated by rodent studies (21,26). The large variability in the skin thickness and the high conductivity of skin can make the results with epidermal montage highly unpredictable. Therefore, placing the electrode over the skull would be a reasonable compromise to mitigate the effects of current shunting by the skin and to improve the focality of stimulation, while still avoiding large E-field peaks as they occur with epidural stimulation.

We did not measure the VE-field at the electrode center with over-the-skull electrodes. The epidural stimulations in Fig. 2 show that the VE-field under the electrode decreases by about a factor of two from the electrode center to 2mm off the edge. If we can extrapolate from over-the-skull stimulations in Fig. 4 at 2mm, the VE-field under the electrode should be about 25mV/mm at the cortical level. The current density at electrode-tissue interface is 56.6 A/m2 for the helical electrode. If we scale the above VE-field value for the current density (142.9 A/m) used by Jackson et al. (27), the result (63 mV/mm) is about 50% higher than the cortical electric fields (42 V/m or 42 mV/mm) predicted in the rat using finite element modeling in their publication. This discrepancy may be due to the differences in the assumed thicknesses and conductivities of the skull and CSF in the model.

### Non-Homogeneity of the Brain

That the skull resistivity was much higher than that of the skin and brain tissue was first demonstrated by early intracerebral voltage measurement studies in human cadavers (28) and anesthetized monkeys (17). Hayes predicted that the high conductivity of the skin and scalp would tend to make the electrical fields more uniform inside the brain. Our data with over-the-skin stimulation does not have the exponentially decaying profile by depth as in over the skull and dura stimulations, and thus agrees with Hayes’ prediction in general. A particular aspect to note is the VE-field peak that occurred consistently at around 1.5-2mm depths in most plots where the gray matter transitions into the white matter and thus a significant change in local conductivity is expected. The lower conductivity of the white matter seems to cause an elevation in the VE-field at the border of the two regions. The fact that the VE-field decline in the gray matter is sharper at 2mm and 4mm horizontal locations than at the electrode center (Fig. 2) must, however, be due to the higher conductivity of the cerebrospinal fluid near the surface, rather than the gray/white matter conductivity differences.

### Electrode Size

The electrode size is the primary factor that can help focalize the induced E-field and the affected volume of tissue (29). Smaller size electrodes require less total current than larger electrodes to induce the same current density at a fixed target point in the brain (30). However the current density at the electrode surface is higher than that of the larger electrodes (31). We did not specifically compare the two different electrode sizes we had due to insufficient statistical power in the data.

### Monopolar vs. Bipolar Montages

Repositioning the return electrode from the shoulder to other muscles around the body introduced changes smaller than 1.36% in the VE-field. This confirms the results reported from finite element models that the voltage profile near the anodic electrode is more or less the same with the monopolar montage so long as the large surface return electrode is placed far enough from the anode. The field steering effect begins to occur when the return electrode is brought near the stimulation electrode, hence approximating a bipolar montage (32,33). Moliadze et al. (34) reported that the stimulation effects can significantly change depending on the distance between the stimulation electrodes, even for the extracephalic placement of the return electrode (e.g. ipsilateral upper arm vs. ipsilateral forearm) in the human. This result is surprising since the current flow patterns should not be affected by the position of the return electrode as long as it is on the same arm. The sensitivity of the results to the location of the return electrode may be higher with epidermal placement of the electrodes due to currents flowing through the highly conductive skin. A practical point raised by our data is that the Ag/AgCl wires deinsulated for several mm at the end and inserted into a muscle can conveniently serve as a return electrode, replacing the large surface transcutaneous electrodes used in other studies (35,36).

That the E-field declines sharply by distance with monopolar montages is advantageous for spatial selectivity if the targeted neural structures are near the brain surface but makes it difficult to achieve significant E-fields at deeper brain regions without causing extreme electric fields near the surface. Attempts to focus the electric field at subcortical brain regions using multiple electrodes will have to deal with this challenge. Horizontal E-field can be maximized using bipolar montages by placing both the cathode and the anode around the head on the sides, if the targeted neural structures are inside the cortical sulci where the somato-dendritic axis of the neurons is oriented horizontally. A recent paper (21) proposed both spatial focusing and time-multiplexing of the stimulus currents via multiple dipoles in order to maximize the horizontal E-field at a focal point inside the brain by taking advantage of the slow time constant of the cellular membranes at subthreshold potentials. While the tES methods enjoy the benefits of being non-invasive, the challenge of focalization may remain as the main disadvantage of the technique in the long run.

### Stimulation Waveform

In our protocol, substituting DC with pulsed stimulation allowed us to overcome the poor DC response of the tungsten recording electrodes. The brain tissue can be treated primarily as a resistive medium hence the recorded amplitudes to be independent of frequency, up to 10kHz and even higher ranges (13,37) owing to the “quasi-static approximation”. Thus, our amplifier’s high cutoff (*f*_*c*_) was set to 10kHz, which has corresponding rise time of 15.9µs *(t*_*r*_*= 1/2 π*_*C*_) for a first-order filter. We took the amplitude measurements 1ms after the pulse transition, allowing several rise times for the signals to stabilize while still being much shorter than the time constant imposed by the lower cutoff frequency of the amplifier (t_r_ = 15.9ms, 10Hz).

A sinusoidal waveform at a constant frequency (e.g. 1kHz) could also be used for the stimulus current in this study instead of rectangular pulses. A stimulus waveform with a single frequency would overcome the bandwidth limitations of the metal recording electrodes and the amplifier, which attenuates the harmonics of a rectangular waveform to different degrees and distorts the recorded waveforms. The advantage of the rectangular waveform, however, is that any mechanical disturbance to the electrode tip during penetration manifests itself not only as a change in the measured amplitudes but also as a distortion in the waveform, and thus alerts the experimenter about the quality of the signals. With sinusoidal waveforms only the signal amplitude would change, which could escape the attention of the experimenter and lead to significant miscalculations of the E-field.

### Comparison with Theoretical Models

Despite the fact that Wiley & Webster equation (25) assumes a homogeneous medium, and does not account for the impedance of the electrode-electrolyte interface and the current redistribution across the electrode surface due to amplitude and frequency dependency of the interface (38), it provides a reference point for comparison. The simplicity of using an analytical equation instead of building complicated models can be much more practical for quick estimations of the E-field underneath a monopolar electrode if that is all that is needed. Comparison of the experimental data with the theoretical model in 2D (Fig. 3) reveals some fundamental similarities but also significant differences. Both plots show that the largest VE-fields occur under the electrode, but the experimental VE-field diminishes within the gray matter for the most part, subsiding to negligible levels in the white matter. In the experimental data the VE-field spreads more in the horizontal direction near the surface most likely due to the high conductivity of the CSF, and perhaps that of the gray matter also to a degree. The theoretical plot assumes a non-conductive medium above the brain and does not account for the presence of the skull. In addition, there are clearly regions of varying conductivities in the experimental data that cause inhomogeneities in the electric field. Lastly, the VE-peaks at the electrode edges that are suggested by the earlier analytical models (25) were not observed in the experimental data. This may be because of the distance (∼0.5mm) allowed between the electrode surface and the brain cortex in our setup, or the spatial smoothing of the E-field peaks may be explained partially by the presence of the electrode-electrolyte interface as predicted by more advanced models (38).

### Practical Considerations

The surface area of the electrode that is in contact with tissue primarily determines the E-field strength in the vicinity of the electrode. Therefore, epidural placement of the stimulation electrode may lead to large variations in the E-fields in the cortex if the electrode moves even by very small amounts. If the electrode contact area with the cortex is well defined, the simple analytical equations (25) can become very useful for predicting the potential and the VE-field underneath the electrode. The equation for the VE-field at the surface (z=0, r=0) is ϕ = −2Vo/(πa). For V_o_ value that gives the best fit with this equation to our data was found to be 58.3mV. Assuming a homogenous medium, an overall conductivity value of 0.57 S/m can be found by substituting this value of Vo into the Newman equation (Vo = J/(4σa)), which is about three times higher than the commonly used conductivity for the brain (0.2 S/m) most likely due to the high conductivity of the CSF (1.65 S/m) (39). For a known electrode diameter and current, one can use this modified conductivity value to estimate the VE-field peak underneath the epidural electrodes. Having an accurate prediction for the vertical E-field near the surface, where most of the cortical neurons are found in an animal with a lissencephalic brain like the rat, can prove to be useful when direct measurements of the field is not possible.

The current distribution across the disk electrodes is predicted to suffer from edge-effects by earlier models (25). Inclusion of the electrode-electrolyte interface into the model produces a more uniform current profile across the surface, although the effect is amplitude and frequency dependent (38). The choice of the electrode material thus is crucial to minimize the electrode-electrolyte impedance and improve reproducibility. In order to prevent the edge-effects from influencing the E- field distribution inside the brain, the stimulation electrode can be kept at a distance from the superficial layers of the cortex. The skull can serve as a spacer in this case. The disk electrode can be attached onto the skull at reproducible stereotaxic coordinates and the top of the electrode can be covered with some nonconductive material to prevent current spreading through the skin. The skull thickness at the point of electrode placement can be standardized to a degree by controlling the weight/age of the animal. Alternatively, one can also make a craniotomy hole and fill it with conductive gel or isotonic saline and position the stimulation electrode at a known distance above the cortex as we did with epidural stimulations. Because normal saline has a conductivity (1.65 S/m) that is about eight times higher than that of the brain (0.2 S/m), the stimulation electrode may electrically be assumed to be at the cortex/saline interface for practical purposes. The highly conductive saline may however be replaced with encapsulation tissue by time in chronic implants. For most reproducible results, the E-field should be measured directly inside the tissue in each animal separately if this can be done without disturbing the intactness of the neurons or the E-field itself.

## V. CONCLUSIONS

This paper provides experimental data as a reference study for more realistic estimates of the electric fields induced in the rat brain during tES studies. The skin attenuates the electric field much more strongly than the skull and causes the current spread more uniformly inside the skull. For focal stimulation, it may be best to place the stimulation electrode on the skull to avoid the skin effect. The electrical field perpendicular to the cortex decreases exponentially near the surface and loses most of its strength within 2mm into the brain underneath the electrode and within 1mm of depth off of the electrode edge. A 100µA current injected through a 1.5mm over-the-skull electrode is predicted to generate ∼25mV/mm at the cortical surface. For epidural placements of the stimulation electrode through a craniotomy hole, a modified value of 0.57 S/m for the brain conductivity can be assumed to estimate the voltage at the cortical surface using the volume conductor equations. Significant E-field peaks occur in the brain parenchyma, most likely due to local conductivity changes, especially at the gray/white matter border. These large fluctuations in the E-field measurements show that the homogeneous volume conductor assumption is too simplistic for modeling the local effects of the injected current.

## ACKNOWLEDGMENTS

Supported by NIH/NINDS Grant No: R21 NS101386.

Author contributions:
Ahmet S. Asan contributed to experimental design, collected the data, and performed signal analysis. Sinan Gok was involved in data collection and drafting the manuscript. Mesut Sahin conceived and designed the experiments, interpreted the data, and reviewed the manuscript.

